# Phylogenetic approaches to identifying fragments of the same gene, with application to the wheat genome

**DOI:** 10.1101/182550

**Authors:** Ivana Piližota, Henning Redestig, Christophe Dessimoz

## Abstract

As the time and cost of sequencing decrease, the number of available genomes and transcriptomes rapidly increases. Yet the quality of the assemblies and the gene annotations varies considerably and often remains poor, affecting downstream analyses. This is particularly true when fragments of the same gene are annotated as distinct genes and consequently wrongly appear as paralogs. In this study, we introduce two novel phylogenetic tests to infer non-overlapping or partially overlapping genes that are in fact parts of the same gene. One approach collapses branches with low bootstrap support and the other computes a likelihood ratio test. We extensively validated these methods by 1) introducing and recovering fragmentation on the bread wheat, *Triticum aestivum* cv. Chinese Spring, chromosome 3B; 2) by applying the methods to the low-quality 3B assembly and validating predictions against the high-quality 3B assembly; and 3) by comparing the performance of the proposed methods to the performance of existing methods, namely Ensembl Compara and ESPRIT. Application of this combination to a draft shotgun assembly of the entire bread wheat genome revealed 1221 pairs of genes which are highly likely to be fragments of the same gene. Our approach demonstrates the power of fine-grained evolutionary inferences across multiple species to improving genome assemblies and annotations. An open source software tool is available at https://github.com/DessimozLab/esprit2.

## Introduction

Thanks to rapid developments in sequencing technology (reviewed in Goodwin et al. 2016), individual laboratories now routinely sequence and assemble entire genomes and transcriptomes. The most well-established short-read sequencing protocols are cost effective and widely applied. However, without reads that are long enough to span repetitive regions, the assembly step remains a challenge with negative consequences on downstream analyses (Lee and Tang 2012; Jiao and Schneeberger 2017).

The challenge of genome assembly is particularly acute in plants, which tend to have large and heavily redundant genomes (Claros et al. 2012; Jiao and Schneeberger 2017). Data from such genomes frequently results in fragmentary assemblies with overestimated gene counts (Denton et al. 2014) and limited utility for downstream purposes such as creation of physical maps used in marker assisted breeding.

One problem in low-quality genome assemblies is that fragments of the same gene can be annotated as distinct entries in genome databases, and thus erroneously appear to be paralogs. However, it is possible to use homologous proteins conserved in other genomes to detect fragments that are likely to be part of the same gene. To our knowledge, four such approaches have been proposed. First, the Ensembl Compara pipeline (Vilella et al. 2009) infers as “gene_split” pairs of apparent paralogs that lie within one megabase on the same strand of the same region of the assembly and do not overlap in the multiple sequence alignment of the family. Restricting these predictions to genes belonging to the same contig greatly reduces the risk of false positive split gene calling, but particularly for fragmented assemblies with many short contigs, this approach detects only a fraction of all splits. Second, ESPRIT (Dessimoz et al. 2011) uses pairwise comparisons to identify non-overlapping pairs of paralogs that have a similar evolutionary distance to homologous sequences in other genomes. The third approach, SWiPS (Li and Copley 2013) is conceptually similar in that it also works based on pairwise alignments—by identifying sets of non-overlapping candidate sequences that have a maximal sum of score with homologous sequences in other genomes. The fourth approach is PEP_scaffolder (Zhu et al. 2016), which relies on high-identity matches of reference proteins to multiple contigs. Thus, like ESPRIT and SWiPS, the approach relies on pairwise alignments. Computationally particularly efficient, it also has the strength of considering a maximum intron length to avoid combining gene fragments that are unrealistically far apart.

Yet for all of these methods, the correct identification of split genes heavily depends on their ability to distinguish fragments of the same gene from fragments from paralogous ones. Ensembl compara and PEP_scaffolder make no attempt to distinguish between the two. As for ESPRIT and SWiPS, although they attempt to identify fragments that match reference proteins consistently—either by requiring consistent evolutionary distances to the reference for all fragments or by requiring consistent best matches for all fragments—these comparisons are inherently limited by the pairwise comparison setting, which loses out on evolutionary information available in a multiple-sequence and tree setting.

Here, to address this problem, we present two complementary phylogenetic methods to identify non-overlapping or slightly overlapping fragments of the same gene that exploit evolutionary relationships across gene families. The first one exploits bootstrap support and the second 2 relies on likelihood ratio tests. We evaluate their performance on an artificially fragmented version of the reference sequence assembly of the wheat chromosome 3B (Choulet et al. 2014). We also compare the two methods, and a meta-approach combining the two methods with ESPRIT, which we call ESPRIT 2.0, to the Ensembl Compara pipeline and ESPRIT 1.0. Finally, we apply new phylogeny-based methods to the early, highly fragmented, draft release of the entire breadwheat genome (International Wheat Genome Sequencing Consortium (IWGSC) 2014) and identify 1221 high-confidence pairs of split genes.

## Materials and Methods

We first introduce our phylogenetic tests of split genes, then proceed to describe the datasets analysed and the evaluation methods. Note that we provide the fine implementations details in the *Supplementary Materials*.

### Phylogenetic tests of split genes

Given a genome assembly with a large number of annotated contigs, the task we face is to figure out which annotated genes actually belong to the same gene, due to annotation mistakes or where the assembler failed to concatenate collinear contigs. Consider therefore two non-overlapping fragments of the same gene sequence. If we perform a multiple sequence alignment of the two fragments together with full-length homologs from other species, and infer a tree based on the alignment, we can expect that the two fragments: i) align to different regions of the multiple sequence alignment (since they are non-overlapping), and ii) sit close to one another in a gene tree inferred from the fragments and the homologous genes.

However, perhaps surprisingly at first sight, although these fragments will generally be close to one another on the tree, they will almost never be inferred as adjacent tips. The reason for this is that since they have no character in common, they cannot be directly compared with each other, only with the other genes in the tree. Thus, there is no information available to infer the distance between them, only to the rest of the tree. The location of the split between the two sequences is therefore undetermined. Furthermore, recall that evolution is modelled as a stochastic process on a tree, with each column in the alignment being a realisation of the process. The two fragments will almost certainly consist of different realisations. Therefore, in the maximum likelihood estimate of the tree, the two fragments’ terminal branches will almost never attach to the exact same place on the tree.

Under the correct model of evolution, however, if the two fragments originate from the same sequence, the difference in the place these are attached to the tree should not be significant.

Here we introduce two tests to infer whether two non-overlapping sequences from the same genome are fragments of the same gene: collapsing branches with low bootstrap support (Efron et al. 1996) and a likelihood ratio test (Wilks 1938).

### Test #1: Collapsing “insignificant” branches

Tree branch support measures are commonly used to gauge the reliability of a branch. Since fragments of the same genes can be expected to be separated by insignificant internal branches on the tree, collapsing branches with low support should result in fragments becoming adjacent tips (also known as “cherries”). Thus, for a given threshold, the test collapses all branches below that threshold and infers as fragments of the same gene all candidates that are cherries.

### Test #2: Likelihood ratio test

The second test to infer fragments of the same gene is a likelihood ratio test (LRT). Our null hypothesis (labelled “*s*” for *split)* is that fragments come from the same gene, and thus can be concatenated into one sequence. The alternative hypothesis (called “*p*” for *paralogs*) is that the two non-overlapping sequences belong to paralogous genes.

*H*_*s*_: *n-1* taxa (split gene)

*H*_*p*_: *n* taxa (paralogous genes)

Test statistic is defined as 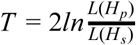, where *L()* denotes the maximum estimator under each hypothesis (Figure 1).

**Figure 1:**
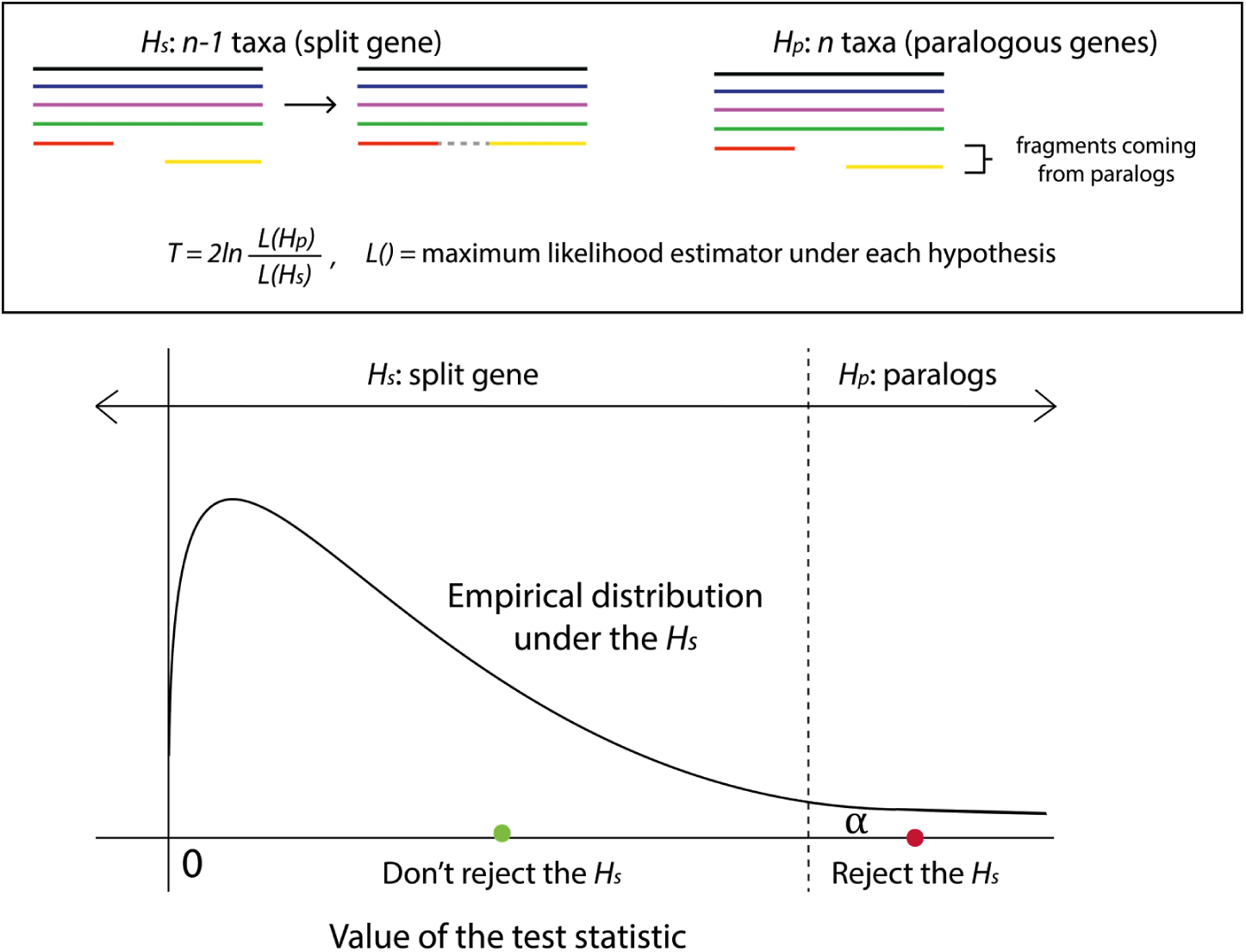
Conceptual overview of the likelihood ratio test. The null hypothesis is that the two fragments come from the same gene (*H*_*s*_) while the alternative hypothesis is that the two fragments come from different paralogous copies (*H*_*p*_). This setup is motivated by the fact that the split gene hypothesis has fewer parameters. However, it is unusual in that failure to reject the test leads to a prediction, and not the other way round.

In a typical setting of the likelihood ratio test, the null model is a special case of the alternative model and the test statistic is chi-square distributed. Since our models are not nested, the distribution of the test statistic given under the assumption that *H*_*s*_ is true is unknown. We can this problem by estimating the empirical distribution under the null using bootstrapping (Efron and Tibshirani 1993; Goldman 1993). Hence, for a particular sample, we:

1. Compute the value of the test statistic; let’s denote it by *T*_*0*_
2. Since we have no prior knowledge on the distribution of the test statistic under the null hypothesis, we estimate the distribution using non-parametric bootstrapping. First, from the multiple sequence alignment used under the *H*_*s*_ we generate *n* artificial alignments of the same length, i.e., *n* bootstrap samples by sampling columns with replacement. Second, we create alignments to be used under the *H*_*p*_ by splitting a target *full-length* gene (i.e. the one made up of two candidate fragments) at the same position as in the original alignment. Finally, we compute the test statistic for each of the *n* samples; let’s denote them by 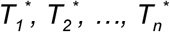. If the sampling is correct, the distribution of 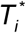, *i = 1, 2,…, n* will converge to the true distribution of the test statistic when *n* → *∞.* Hence, if repeated many times, the distribution of the bootstrap sample test statistic values will approximate the distribution of the unknown test statistic. Throughout this project we set *n* to 100 unless otherwise stated.
3. Compute bootstrap *p*-value as the proportion of samples with likelihood equal or above that of the input data: 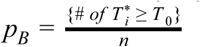

## Implementation of the tests

As input candidate pairs, we identify, among all the protein sequences of a target genome, those that belong to the same gene family—either established by Ensembl Compara or defined as deepest hierarchical orthologous groups as inferred by OMA (Altenhoff et al. 2013)). We further require that fragments be non-overlapping (less than 10% positions in the same alignment column, using Mafft v7.164b (Katoh and Standley 2013)).

The LRT requires computing maximum likelihood estimates, i.e. finding an optimal tree under both *H*_*s*_ and *H*_*p*_. Under the *H*_*s*_ hypothesis, fragments are part of the same gene. Hence, in order to find a maximum-likelihood tree under the *H*_*s*_, we concatenate the candidate fragments into a single sequence. To correct for some cases when a tree-building method gives a suboptimal tree, which may result in the estimated *T*_*0*_ < *0* (impossible in theory), we performed two tree as adjacent tips), and proceeded with the tree with higher likelihood.

Some genes might be involved in multiple predictions, i.e., in more than one pair of fragments coming from a split gene. If all these multiple predictions span different parts of the sequences, we conclude that the gene is split in more than two pieces and consider these predictions as non-ambiguous. If by contrast more than one prediction spans over a common part of the sequence (which might be the case if the fragments come from very closely related paralogs, or if alternative splicing variants of the same gene are erroneously annotated as separate genes), we report the overlapping predictions as ambiguous.

## Datasets and evaluation methodology

As a test case for evaluation and application of the methods, we used proteome of bread wheat, i.e., *Triticum aestivum* cv. Chinese Spring. In 2014, the International Wheat Genome Sequencing Consortium (IWGSC) published a highly fragmented chromosome-by-chromosome survey sequence of the bread wheat genome (International Wheat Genome Sequencing Consortium (IWGSC) 2014). The same year, Choulet *et al.* (2014) published a high-quality reference sequence of bread wheat chromosome 3B. The two provide a good basis to evaluate our methodology on a challenging dataset.

As customary in the field, we determine the quality of the methods by measuring the precision and recall. Here the recall measures the proportion of fragmented genes that the methods can identify. The precision penalises for erroneous predictions by measuring the proportion of predictions that are indeed fragmented genes. For both measures, we simulated fragmentation on the wheat 3B reference sequence. In a subsequent experiment, we applied the tests to the wheat 3B survey sequence and validated predictions using the wheat 3B reference sequence. In this case the total number of fragmented genes is unknown, so we could only count the number of correct and wrong predictions, and calculate the precision.

Finally, we applied the methods to the rest of the wheat survey sequence to infer split genes in bread wheat proteome.

### Random fragmentation of the wheat 3B reference assembly (recall)

To determine the recall of the methods, we simulated fragmentation on genes assigned to a high-quality assembly of bread wheat chromosome 3B (3B reference sequence). All genes and their gene families were obtained from Ensembl Plants, release 31. We randomly chose one hundred genes, each at least 100 amino-acids long, and split them at a random position such that both fragments are at least 50 amino-acids long. All alignments were performed using Mafft v7.164b with default parameters. Gene trees were build by FastTree v2.1.8 (Price et al. 2010), also with a default set of parameters.

In addition, we simulated fragmentation in a more challenging setting, i.e. on small gene families typically containing only evolutionarily very close paralogs. As a source of homologous groups, we used hierarchical orthologous groups (HOGs). They were computed by the GETHOGs algorithm with a default set of parameters on the input dataset comprised of thirteen plants: bread wheat and twelve flowering plants exported from OMA Browser (Altenhoff et al. 2014) (Suppl. table 1).

### Introducing non-overlapping paralogs in wheat 3B reference assembly (precision)

To inspect cases where the methods incorrectly predict split genes, we simulated fragments from pairs of paralogs assigned to the bread wheat 3B reference sequence using the same datasets as above. We chose one hundred pairs of same-species paralogs, cut them at a random position and took two complementary fragments (one from each initial gene) each being at least 50 amino-acids long. Again, MSAs were obtained by Mafft v7.164b (default parameters) and gene trees by FastTree v2.1.8 (default parameters).

Similarly as above, we also simulated more challenging cases of fragmentation. We used the same set of HOGs as in the previous section.

### Validation on 3B survey assembly

To assess predictions on the real data containing fragmented genes, we applied our approaches to a low-quality assembly of bread wheat chromosome 3B - 3B survey sequence (IWGSP1; 2013-11-MIPS), and compared the predictions with the high-quality assembly of chromosome 3B (“3B reference sequence”) downloaded from URGI (https://urgi.versailles.inra.fr). As gold standard, we mapped sequences between the two assemblies using BLAST+ v2.2.30 (Camacho et al. 2009).

For the predictions, we used the same reference species as in the simulations on HOGs (see previous two sections) which we again exported from OMA Browser (Suppl. Table 2^1^). We computed gene families by GETHOGs algorithm with a default set of parameters. We generated 500 bootstrap samples for each family and performed both tests on fragments overlapping less than 10%. Sequences were aligned with Mafft v7.164b (default parameters) and trees built with FastTree v2.1.8 (default parameters) as above. In addition, we also computed HOGs with a different set of parameters and repeated the rest of experiment.

For the assessment, the mapping of sequences between the survey and high-quality genomes was not straightforward because the two differ not only in the degree of fragmentation, but also in some of the sequences themselves due to sequencing error, contamination etc. To allow for a bit of tolerance while still maintaining unambiguous mapping between the two, we required hits to cover at least 95% of the corresponding query, the percentage identity in these matching regions to be at least 95%, and the hit to be unambiguous. As a stringent control, we also performed a validation where, in addition to these two requirements, we only allowed mismatches to occur at the ends of a query sequence.

### Comparison to established methods and meta-approach

As a point of comparison, we employed Ensembl Compara pipeline and ESPRIT on the same 3B survey sequence as above. Again, the obtained predictions from each method were mapped to the 3B reference sequence by BLAST+ v2.2.30 to inspect if predicted pairs belong to the same gene or not, requiring both coverage and percentage identity to be at least 95%. Validated predictions were compared to the results from Validation experiment on 3B survey sequence with the same BLAST+ criteria.

To obtain a comparable set of predictions on the 3B survey sequence using public results available from the Ensembl Compara pipeline, we filtered “gene_split” pairs from their homologies file (release 21). We took only pairs where both genes were at least 50 amino-acids long and such that, when corresponding gene family was aligned with Mafft v7.164b, candidate genes overlapped for less than 10%. We also included cases where more than two genes were inferred as a part of the same gene given that no two genes involved overlapped for 10% or more. Since some of the sequences could not be found in the OMA Browser dataset used for validating Collapsing and LRT approach, we classified Ensembl predictions into two groups: those that could be found in the OMA Browser dataset, and hence, included in the comparison, and those that could not.

Another set of predictions was obtained by running ESPRIT on the same 3B survey sequence data using twelve reference plants (the same dataset as in the Validation section, Suppl. table 2) keeping all parameters default but increasing the required length of the candidate genes to be at least 50 amino-acids (`MinSeqLenContig:= 50`). We only considered a confident unambiguous set of predictions (`hits.txt` file).

In addition, we considered a meta-approach ESPRIT 2.0 - encompassing ESPRIT and the new combined approach. It takes the union of predictions made by ESPRIT and our joint method (Collapsing branches with support lower than 0.95 and likelihood ratio test with significance of 0.01).

### Inferring split genes on the rest of the wheat survey assembly

Finally, we employed the tests to infer fragmented genes in the first draft release of the predicted genes in whole bread wheat genome, i.e., *Triticum aestivum* cv. Chinese Spring proteome (IWGSP1; 2013-11-MIPS). We considered only candidate fragments assigned to the same chromosome and the same chromosome arm. We used the same reference genomes as in the previous analyses with HOGs (see above). Based on simulations and validation on 3B survey sequence, we determined a set of parameters used for predictions. In particular, we ran GETHOGs with default parameters and allow candidate fragments to mutually overlap less than 10% in the corresponding MSA. We used Mafft v7.164b to get alignments and FastTree v2.1.8 to construct trees, both with their default set of parameters. Finally, we chose 0.95 as a threshold for collapsing and set significance of the LRT to 0.01.

## Results

Recall that we aim to identify fragments of the same gene wrongly annotated as separate genes in a genome of interest, leveraging genomes of related species. In the previous section, we introduced two phylogenetic methods: one based on collapsing branches with low bootstrap support and the other relying on a likelihood ratio test (LRT). To evaluate the methods and determine parameters for predictions on the bread wheat assembly, we took two approaches. First, we simulated fragmentation on the real data to calculate recall and precision. Then we applied both methods to the bread wheat chromosome 3B survey sequence and validated predictions with respect to the 3B reference sequence. Finally, based on the best parameters obtained from these analyses, we applied the method to infer split genes in the 20 other chromosomes of the survey wheat assembly.

### Artificial fragmentation of the wheat 3B reference assembly

To assess our methods, we first simulated fragmentation in 100 protein sequences from the high-quality wheat 3B reference assembly and tried to recover these pairs. Our simulations also included one hundred pairs of non-overlapping fragments generated from pairs of randomly selected paralogous genes—which can be very difficult negative cases if the paralogs are near-identical.

On these challenging simulations, the collapsing test yielded high precision (0.85-0.88) and moderate recall (0.20-0.58), while the LRT performed the other way round, yielding moderate precision (0.56-0.64) and high recall (0.81-0.99) (Fig. 2a, Suppl. File S1, Suppl. File esprit2_simulations.tar.gz).

**Figure 2.**
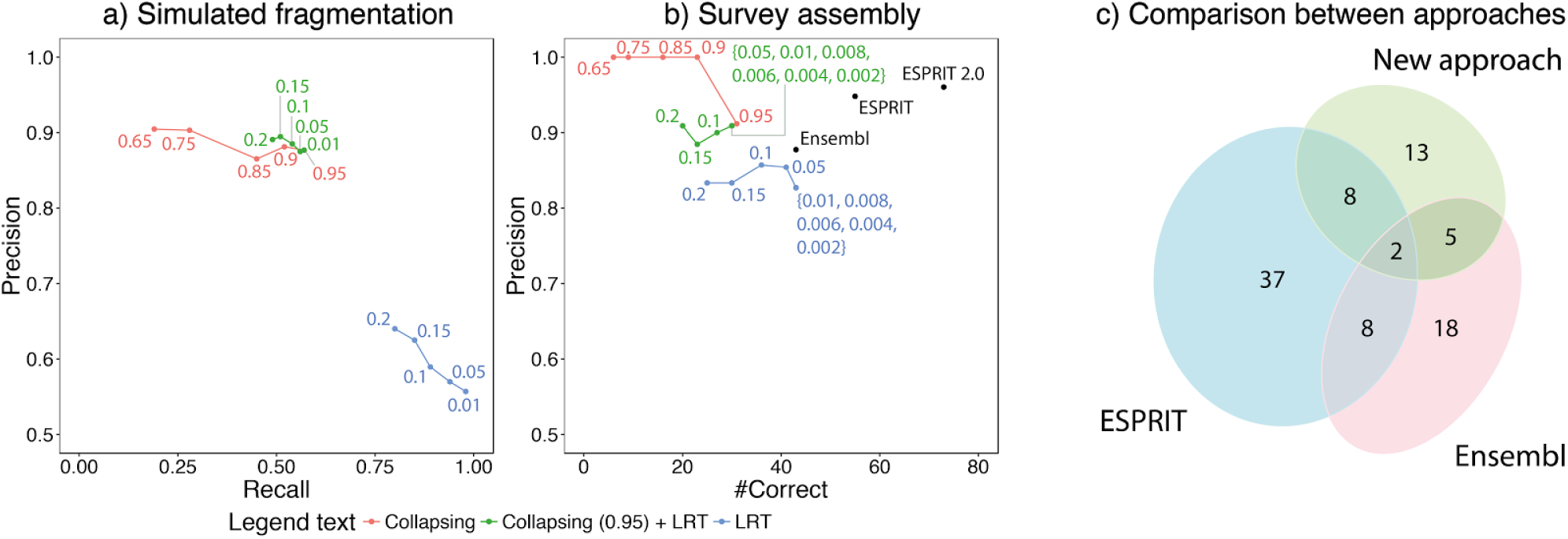
Evaluation of the methods. a) Wheat genes from the high-quality wheat 3B chromosome were artificially fragmented and recovered by the collapsing, likelihood ratio test (LRT), and a combination between the two. b) Split genes inferred on the low-quality (“survey”) wheat genome were validated using the high-quality wheat 3B, and comparison with three other approaches (Ensembl Compara, ESPRIT and ESPRIT 2.0). ESPRIT 2.0 combines ESPRIT’s and the predictions inferred when combining collapsing approach (threshold 0.95) and LRT (significance 0.01). c) The number of predictions on 3B survey sequence classified as correct in the BLAST+ validation. “New approach” denotes a combination of collapsing approach (threshold 0.95) and LRT (significance 0.01)

We also evaluated an approach that combines our two methods. A split gene was inferred if both methods were in agreement. This approach resembled the recall and precision of the collapsing approach (with the same threshold) but with slightly higher precision (Fig. 2a, Suppl. File S1).

As a control, we performed another set of simulations using a different set of input homologous sequences—OMA hierarchical orthologous groups (HOGs) containing protein sequences from thirteen plants including wheat (Suppl. File S1, Suppl. File esprit2_simulations.tar.gz). Precision of the collapsing test was again high (0.73-0.81) while recall varied between 0.30 and 0.78. Precision of the LRT was moderate to high (0.51-0.89) and the recall was high (0.70-0.75) (Suppl. fig. 3a).

### Validation on 3B survey assembly

To further assess the tests and identify suitable parameters, we applied our method on the chromosome 3B of the bread wheat survey genome(International Wheat Genome Sequencing Consortium (IWGSC) 2014); indeed, this is the one chromosome arm for which a high-quality referencewas available (Choulet et al. 2014) but which was not used for creating the draft whole-genome assembly.

Overall, the methods achieved higher precision than when applied to simulated fragmentation (Fig. 2b). The analysis showed particularly high precision of the collapsing approach. The absolute recall rate could not be easily assessed on these real data; instead, we considered the number of correctly predicted HOG annotations as a surrogate for recall, yielding results highly consistent with the simulations (Fig. 2b).

One challenge with this setup was the fact that the draft survey sequence assembly contains other types of problems, such as sequencing errors or ~10% contamination from other chromosomes (International Wheat Genome Sequencing Consortium (IWGSC) 2014)). If we only consider fragments that can be perfectly mapped between the draft whole-genome assembly and the reference assembly (no mismatch in their central part, see *Supplementary Materials*), the number of predictions that could be validated diminishes, but on the remaining set, our approaches showed even higher precision (Suppl. fig. 3c and 3d), indicating that the performance reported in Fig. 2b is conservative.

Control experiments also gave consistent results (Suppl. File S2, Suppl. File esprit2_validation.tar.gz). As expected, relaxing parameters yielded more predicted split genes, but at a cost of lower precision (Fig. 2b vs. Suppl. fig. 3b).

### Comparison to established methods and meta-approach

To gain further insights into the performance of the proposed approaches, we compared them to two existing methods, namely Ensembl Compara pipeline (which however cannot easily run on custom genome data) and ESPRIT, as described in the Methods. Both methods were applied to the 3B survey sequence and then validated against the 3B reference sequence using BLAST+ (Suppl. File esprit2_comparison.tar.gz). We also considered a meta-approach, which we call ESPRIT 2.0, comprising ESPRIT and a combination of the collapsing approach (threshold 0.95) and LRT (significance 0.01).

In terms of the number of correct predictions, Ensembl Compara and ESPRIT performed equally well or better than our approaches displaying high precision (Fig. 2b and Suppl. table 3). Further analysis showed that predictions from different methods are rather complementary and worthwhile taking into account (Suppl. fig. 4). Hence, the meta-approach, ESPRIT 2.0, inferred by far the biggest number of correct predictions with high precision (Fig. 2b and Suppl. table 3).

### Predictions on the rest of the survey assembly

Finally, we applied our tests to infer split genes on the rest of the bread wheat genome, i.e., all chromosomes other than 3B. Based on the analyses on simulated fragmentation and between two assemblies (see above), we determined parameters for the tests. For each chromosome arm, we obtained gene families by running OMA GETHOGs with default parameters. In the collapsing approach, we collapsed all branches with bootstrap support less than 0.95, and we performed the likelihood ratio test with the significance level of 0.01. The intersection of predictions identified 1442 pairs in total: 1221 unambiguous and 221 ambiguous cases. The distribution of the number of predictions per chromosome is shown in Fig. 3 (see also Suppl. File S3) while fragment IDs are provided in Suppl. file esprit2_predictions_wheat.tar.gz.

**Figure 3.**
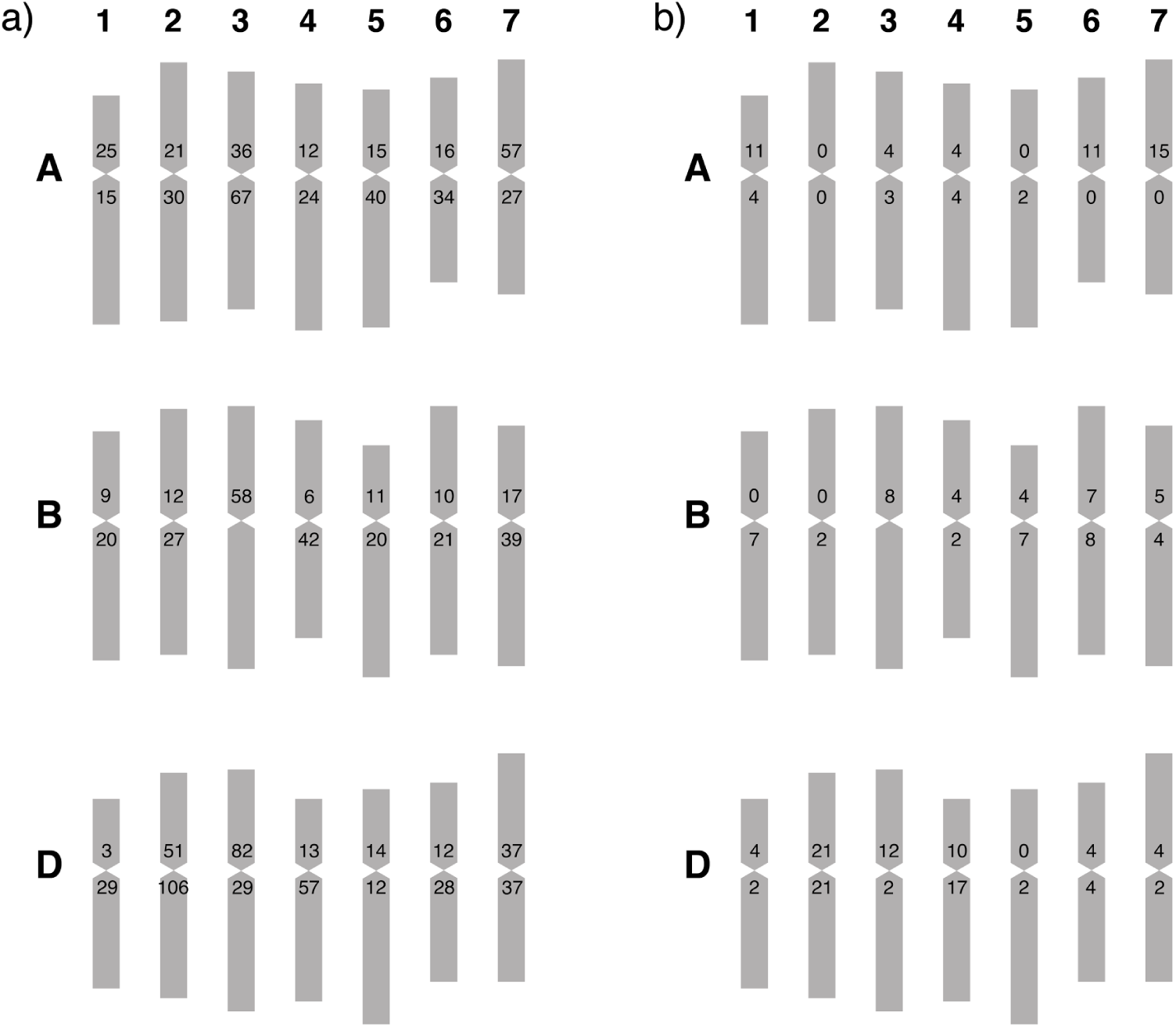
High-confidence inferred gene splits on the wheat genome. a) Number of unambiguous predictions for each chromosome arm. b) Number of ambiguous predictions (i.e. for which there is more than two candidate fragment for a single juncture). Pairs of fragments are inferred separately for each chromosome arm of flow-sorted *Triticum aestivum* cv. Chinese Spring, except chromosome 3B, for which the analysis was performed on the entire chromosome.

## Discussion and outlook

Despite technological and algorithmic advances, genome assembly and annotation remains a challenge, especially for large polyploid genomes with complex evolutionary histories. Genes often remain fragmented and fragments get annotated as separate genes. Our work demonstrates that using available assemblies of related species can provide enough information to recognise some of those cases and obtain full-length genes.

We developed two approaches and showcase their good performance on a challenging proteome of hexaploid bread wheat (*Triticum aestivum* cv. Chinese Spring). In simulations and validation, both of which were performed on the real data taking into account all its complexities, an approach relying on collapsing gene tree branches showed lower recall and higher precision than a likelihood ratio test (Fig. 2). As a trade-off between precision and recall, we propose taking an intersection of their predictions, as we did in the quest for fragmented genes in the wheat survey sequence dataset. As our stringent simulation and real data assessment shows, the inferred split genes are highly specific. The performance is even better when we combine the new phylogeny-based tests our earlier pairwise approach “ESPRIT”.

The two main inherent challenges of *in-silico* split gene inference are the confounding effect of close paralogs and the variation in the rate of evolution along the sequences. Indeed, sometimes fragments come from identical or nearly identical paralogs and there is not enough information to distinguish fragments belonging to one gene from another. Hence, we are more likely to make a false positive prediction (Suppl. Fig. 5). As for evolutionary rate heterogeneity across the protein length, this can pose problem because fragments of the same genes can wrongly appear to be coming from distinct sequences. Consider for instance a protein composed of two domains—one slowly evolving and one fast evolving. If we consider each domain as a distinct sequence and look at their position in a gene tree including full-length homologous counterparts, the branch lengths connecting these fragments to the rest of the tree may have markedly different lengths. As a consequence, the increase in likelihood obtained by having distinct branches for each fragment may be sufficiently large for our test to erroneously infer that the fragments come from distinct sequences (see Suppl. Table 4 for an actual example). It may be possible to address this problem by more explicitly modelling variation of rate among sites.

At a practical level, predictions heavily depend on the choice of two parameters: a threshold for collapsing branches and a significance level for likelihood ratio test. Lower, more stringent thresholds for collapsing yield more confident predictions, while higher, less conserved thresholds will produce more predictions but less confident. Similarly, a higher significance of the likelihood ratio test will result with less but more confident predictions. Obtaining more predictions can be achieved by lowering the significance of the test at the cost of their lower confidence. Overall, it is important to choose thresholds depending on the application. For example, a higher number of predictions can be favourable for comparison with other data.

Predictions also depend on the input families. Bigger gene families facilitate more predictions (Fig. 2, Suppl. fig. 3) but also result in more ambiguous calls, i.e., cases where a fragment is involved in multiple predictions (Suppl. File S2). We observed fewer false positive predictions when we simulated fragmentation on bigger gene families where we were more likely to randomly split a pair of more distant paralogs in comparison to small gene families which are more likely to contain only very close paralogs (Fig. 2a, Suppl. fig. 3a). However, the results of validation indicate that the methods are still able to identify a reasonable number of split genes with high precision even when small gene families are used.

Throughout this project, we fixed some of the parameters. First, we considered only genes at least 50 amino-acids long. Shorter sequences contain less information thus make phylogeny reconstruction more challenging; at the same time, the benefit of putting together short fragments is also more limited. Second, we required candidate fragments to overlap less than 10%. Increasing the overlap increases the number of candidate pairs and, consequently, the number of predictions including false positive and ambiguous predictions. Finally, we used Mafft v7.164b to align gene families and FastTree v2.1.8 to reconstruct gene trees, both with their default parameters due to their convenience and speed. Exploring their parameter space or using more suitable tools for the dataset of interest could contribute to higher precision and recall.

As often with new approaches, the likelihood ratio test still has room for improvement. Currently, we compute the distribution of the test statistic empirically, via resampling. We computed up to five hundred samples per test which, given the simulations and validation, seems to be enough here; yet the convergence of the distribution could be explored. Increasing the number of samples might lead to significantly better approximation of the distribution and more accurate results. In addition, parameterising the distribution of the test statistic would reduce computational time and memory usage.

Since both tests rely on evolutionary relationships, some of the mistakes could be avoided by implementing more realistic evolutionary model. This is of particular importance for cases which are missed due to differences in evolutionary rates across the length of the gene.

To further improve the performance, one could try to find optimal parameters for the dataset of interest and application in question. Different strategies could be used to obtain input families as well as alternative tools for alignments and methods with more exhaustive optimal tree search.

But already in its present form, as the large number of detected split genes in the wheat genome illustrates, our approach is already proving highly useful. All computer code is available for reuse as a user-friendly package (https://github.com/DessimozLab/esprit2) that we hope will help make phylogeny-based detection of split genes a routine step in genome assembly and annotation.

## Acknowledgements

We thank Adrian Altenhoff for help packaging DLIGHT 2.0 as a standalone software tool. Computations were performed on the University College London Computer Science cluster. We gratefully acknowledge funding from Bayer CropScience NV, University College London, the Swiss National Science Foundation (grant PP00P3_150654 to CD) and the UK Biotechnology and Biological Sciences Research Council (grant BB/L018241/1 to CD).

## Supplementary Materials

### Implementation of the tests

#### Input data

##### Concatenating non-overlapping fragments into a single sequence

Concatenation was done at the level of multiple sequence alignment, i.e., we aligned a gene family and then replaced the two fragments with a newly created sequence containing residues from both fragments and gaps at the remaining positions.

##### Working with overlapping fragments

Given a chosen threshold *L* ∈ *< 0, 1>*, two candidate fragments were allowed to overlap for less than *L*100%* of the region length each of them spanned in the multiple sequence alignment. Looking at the alignment from left to right, this means that the end of one fragment and the beginning of the other overlap. In order to concatenate them, we first determined the *middle* of the overlap and edited the sequences as follows. In the sequence on the left, we kept all positions the same up to the *middle* and replaced the remaining residues with X’s. Similarly, in the sequence on the right, we edited the beginning of the sequence by replacing all residues up to the *middle* with X’s and then kept all remaining residues as they were. This way we got two fragments with non-overlapping known residues in the alignment. They were used as such in both tests. For trees under the *H*_*s*_ model in the likelihood ratio test, these modified sequences were concatenated as described in the section on concatenating non-overlapping fragments (see above).

##### Input topology

As an input topology in the search for maximum likelihood tree under the *H*_*p*_ model, we used a modified tree from the *H*_*s*_ model. We bifurcated a node (leaf) with the candidate gene and set new branches’ lengths to zero. We set the support of a branch leading to fragments’ parental node to 0.5 (Suppl. fig. 1).

**Supplementary figure 1:**
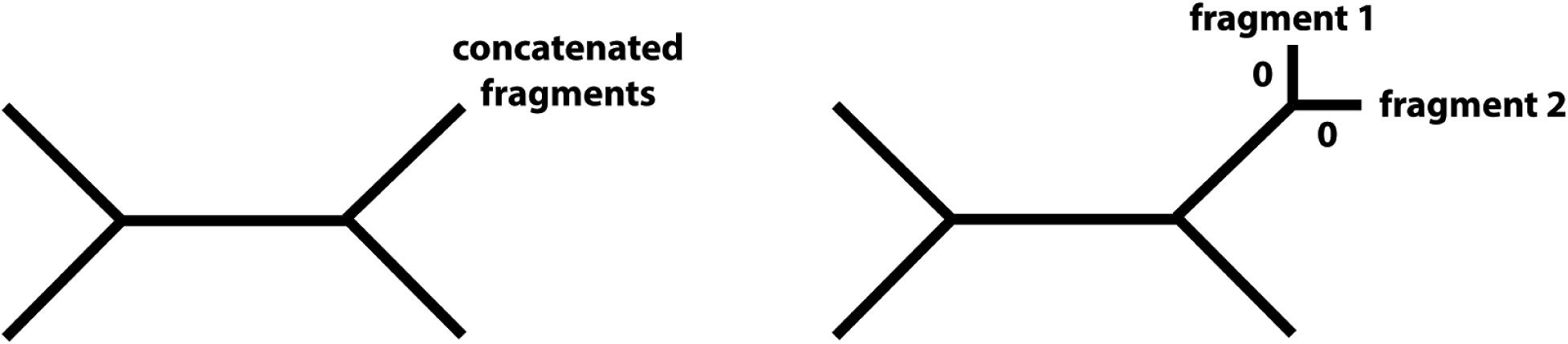
Input topology for the likelihood ratio test. Left: Maximum likelihood tree under the *H*_*s*_. Right: Modified tree to be used as an input tree in the *H*_*p*_ model.

### Datasets and evaluation methodology

#### Random fragmentation of the wheat 3B reference assembly (recall)

##### Fragmentation

First, we aligned corresponding gene families using Mafft v7.164b with default settings. Then we chose a random position *n* in the alignment such that the target gene, i.e., the one we want to fragment, contains at least 50 amino-acids in the first *n* positions of the alignment and at least 50 amino-acids right of the chosen position. First *n* positions of the aligned target gene extended by *alignment_length-n* gaps form one fragment, while the second fragment is formed from *n* gaps extended by the rest of aligned target sequence. The original target sequence is then replaced with newly formed fragments while the rest of the gene family is kept the same (Suppl. fig. 2a).

##### Simulations on HOGs

We performed the experiment as follows:

1) We computed HOGs (input data in Suppl. table 1) using GETHOGs algorithm implemented in OMA standalone keeping default settings.

2) A random 3B gene was selected, and a top-level HOG containing the gene was aligned using Mafft v7.164b (default settings). The gene was split at a random position (see above).

3) All necessary trees were computed using default settings in FastTree v2.1.8.

4) Both methods were applied to all pairs of candidates.

#### Introducing non-overlapping paralogs in wheat 3B reference assembly (precision)

##### Fragmentation

Again, we aligned gene families using Mafft v7.164b with default settings. Genes from a randomly chosen pair of paralogs were assigned to *sequence 1* and *sequence 2* at random. Then we chose a random *n* such that the first *n* positions of *sequence 1* and last *alignment_length-n* positions of *sequence 2* each contain at least 50 amino-acids. These two subsequences form the basis of simulated fragments, one extended by gaps on its right end and the other extended by gaps at its left end (Suppl. fig. 2b). If there was no such *n*, the pair was discarded.

##### Simulations on HOGs

Analogous to the *Simulations on HOGs* in *Random fragmentation of the wheat 3B reference assembly (recall)*.

**Supplementary figure 2:**
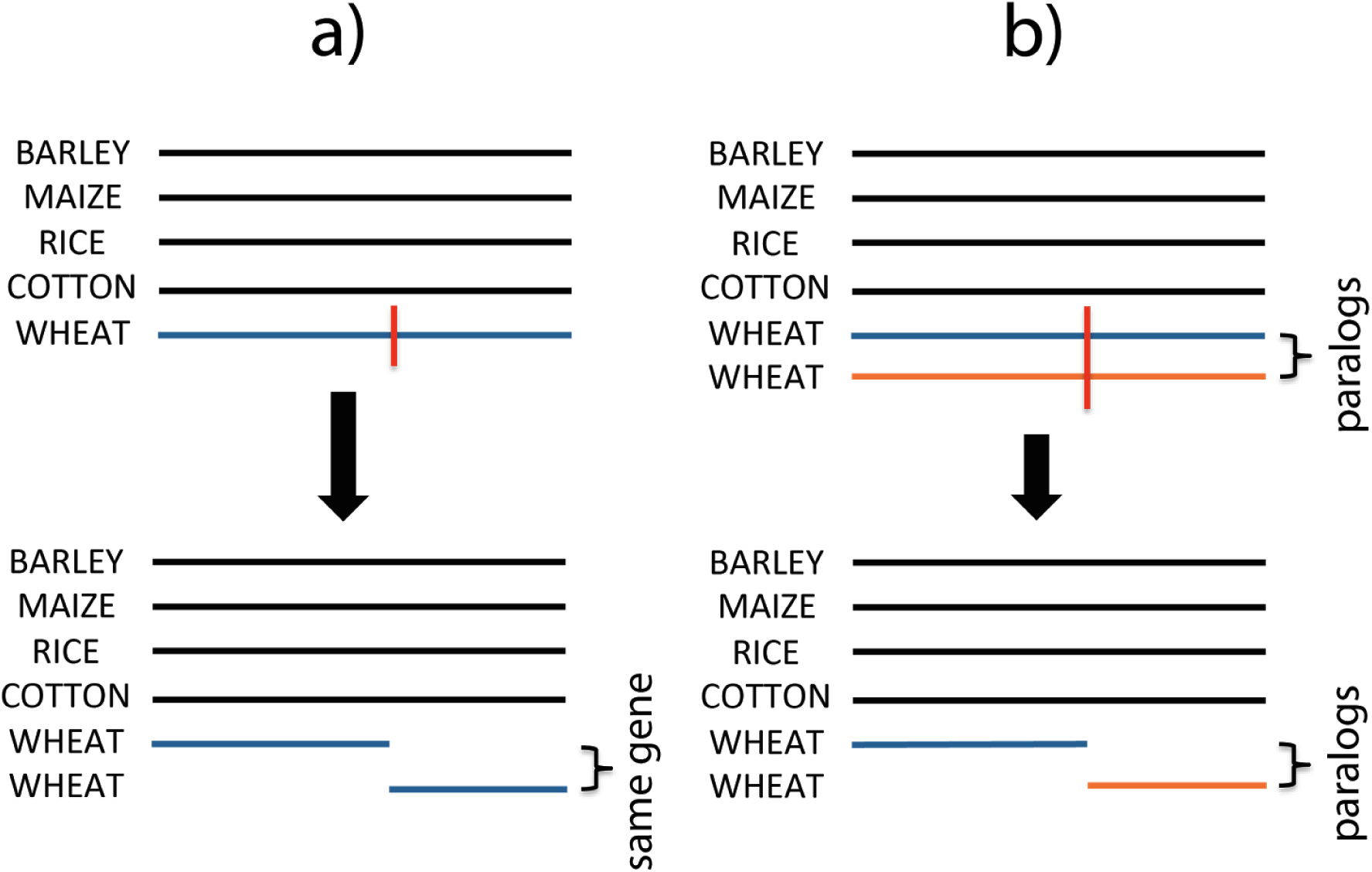
Simulating fragmentation. a) Simulating fragments coming from the same gene, b) Simulating fragments coming from paralogs.

**Supplementary table 1:** Proteomes exported from OMA Browser and used as input data for GETHOGs algorithm in simulations. The second column contains information on the database release that OMA browser retrieved an assembly and annotation from.

| Species | Database |
| --- | --- |
| Aegilops tauschii | Ensembl Plants 21 |
| Arabidopsis thaliana | Ensembl Plants 20 |
| Brachypodium distachyon | Ensembl Plants 21 |
| Hordeum vulgare var. distichum | Ensembl Plants 16 |
| Oryza brachyantha | Ensembl Plants 21 |
| Oryza glaberrima | Ensembl Plants 21 |
| Oryza sativa subsp. indica | Ensembl Plants 21 |
| Oryza sativa subsp. japonica | Ensembl Plants v7 |
| Setaria italica | Ensembl Plants 21 |
| Sorghum bicolor | Sbi1_4 |
| Triticum aestivum cv. Chinese Spring | Ensembl Plants 26 |
| Triticum urartu | Ensembl Plants 19 |
| Zea mays | Ensembl Plants v8 |

#### Validation on 3B survey assembly

**Supplementary table 2:**
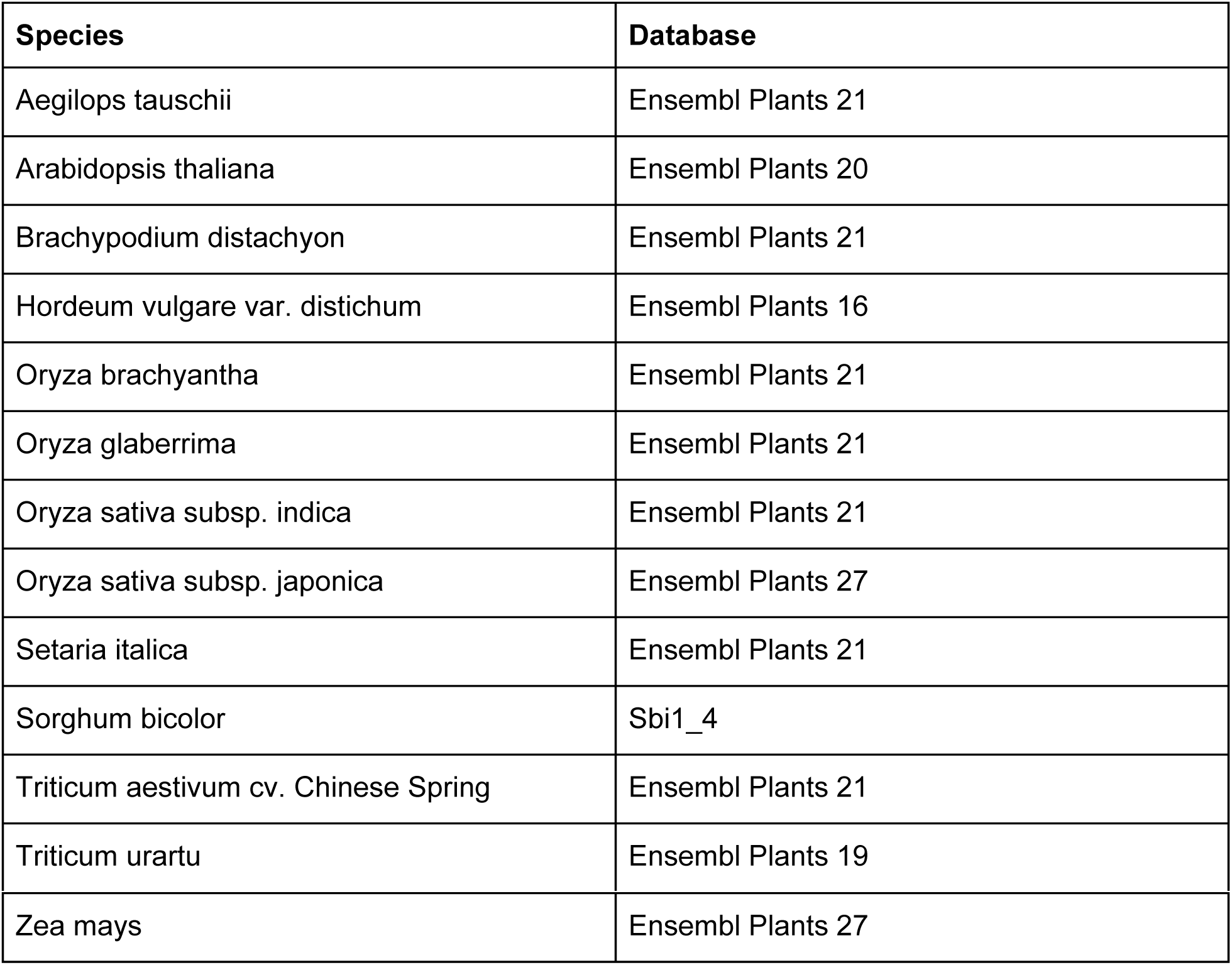
Proteomes exported from OMA Browser and used as input data for GETHOGs algorithm in validation on *Triticum aestivum* cv. Chinese Spring chromosome 3B. The second column contains information on the database release that OMA Browser retrieved an assembly and annotation from.

##### Procedure

We performed the experiment as follows:

1) We computed HOGs using GETHOGs algorithm implemented in OMA standalone keeping default settings. Since OMA requires sequences to be longer than a certain threshold, in order to consider shorter sequences in the target wheat genome, we renamed WEAT.fa file in OMA/Cache to WHEAT.contig.fa.

2) Each top-level HOG was aligned using Mafft v7.164b (default settings) and searched for candidate fragments. Two fragments were called a pair of candidates if their overlaps were below 0.1, i.e., 10%.

3) All necessary trees were computed using default settings in FastTree v2.1.8.

4) Both tests were done on all pairs of candidates.

##### Control

Since GETHOGs algorithm was not developed for the purpose of surveying genome assemblies, its default parameters might not be optimal for this purpose. In particular, a set of default parameters might be too conservative so we repeated the experiment described above but with some parameters lower than default (`MinScore:= 150, LengthTol:= 0.4,ReachabilityCutoff:= 0.3`). This yielded bigger HOGs and hence more candidates to test.

##### BLAST+ validation

We developed two modes of BLAST+ validation. A less stringent one conditions only on query coverage per hit (`qcovs`) and %identical matches (`pident`). A more stringent one allows mismatches to be only at the ends of a query sequence.

Less stringent validation

For each unambiguous or ambiguous prediction we:

1. Take initial non-modified sequences of both fragments and BLAST+ them against high-quality assembly of chromosome 3B (`-evalue 0.001`)
2. For each query sequence, identify BLAST+ hit(s) with the highest bitscore (`bitscore`). Keep only hit(s) with `qcovs` >= 95 and `pident` >= 95, if any. If there are no such hits for any of the queries, the pair cannot be validated.
3. For each query and its hits from 2., keep only hits with the highest `qcovs.` If there are multiple hits per query that satisfy the criteria, then filter out all hits with `pident` lower than the highest present. If there are still multiple hits for any of the queries, we consider that an ambiguous mapping and do not validate the pair.
4. If both queries have the same best hit, the prediction is considered to be correct. Otherwise, we consider it wrong.

##### More stringent validation

All steps are the same as in *Less stringent validation* except the step 2. Here, in addition to `qcovs` >= 95 and `pident` >= 95, we require all mismatches between a query and a hit to be at the ends of a query sequence.

Let’s say that our tolerance length is M. Suppose that first N_1_ and last N_2_ positions of a query are not covered by a hit. If N_1_ > M or N_2_ > M, then the hit does not pass the criteria. For a given query and a hit such that 0 < = N_1_, N_2_ < = M, consider their BLAST+ alignment. We allow mismatches to be only in the query’s first M-N_1_ or last M-N_2_ aligned positions, and we set M=5.

### Results

**Supplementary figure 3:**
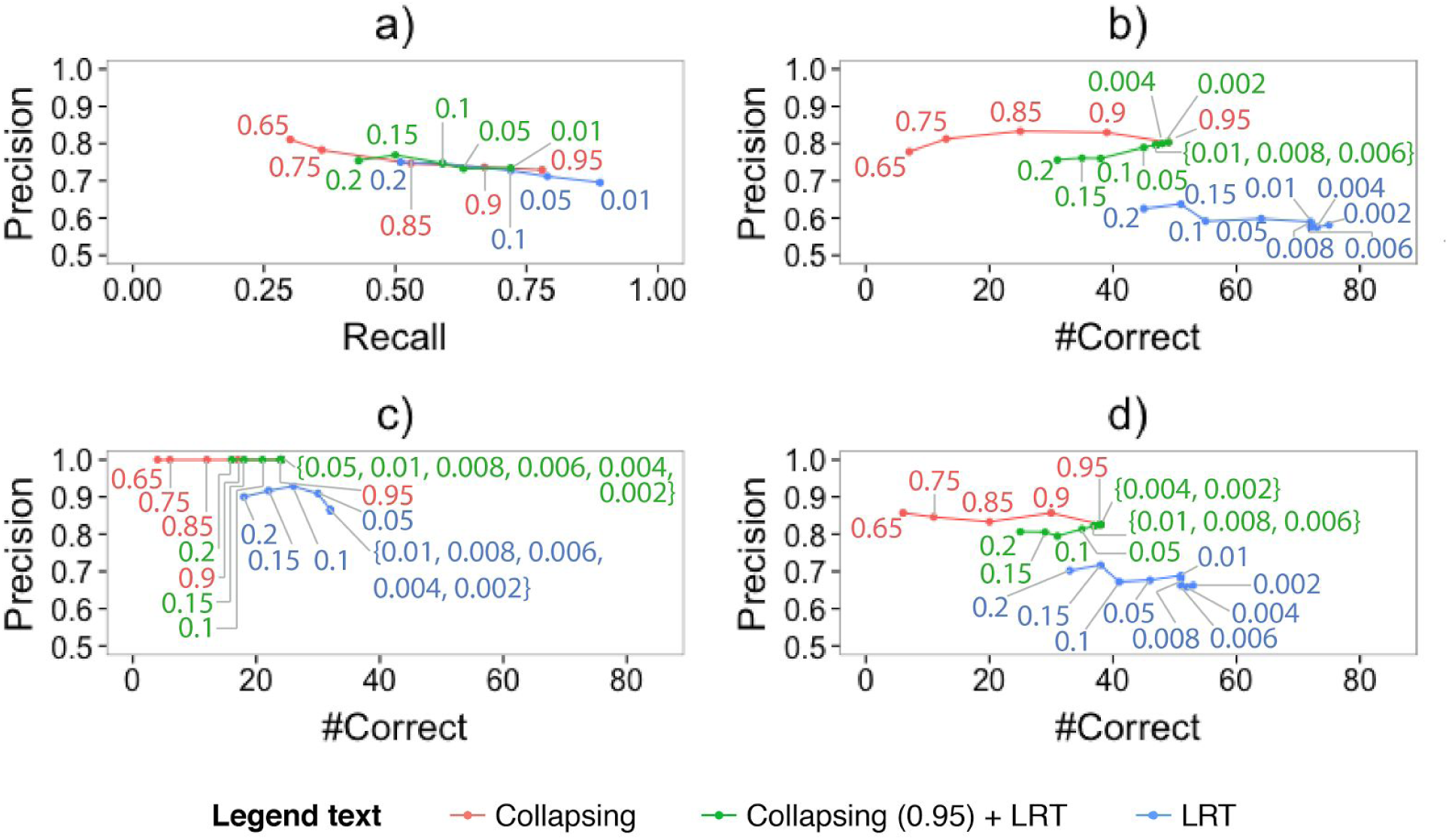
Results of control experiments: simulations and validations. a) Simulated fragmentation on HOGs with default settings in GETHOGs algorithm, b) Validation on 3B survey sequence using HOGs with relaxed parameters in GETHOGs algorithm, less stringent BLAST+ validation, c) Validation on 3B survey sequence using HOGs with default settings in GETHOGs algorithm, more stringent BLAST+ validation, d) Validation on 3B survey sequence using HOGs with relaxed parameters in GETHOGs algorithm, more stringent BLAST+ validation

### Comparison to other methods

**Supplementary table 3:**
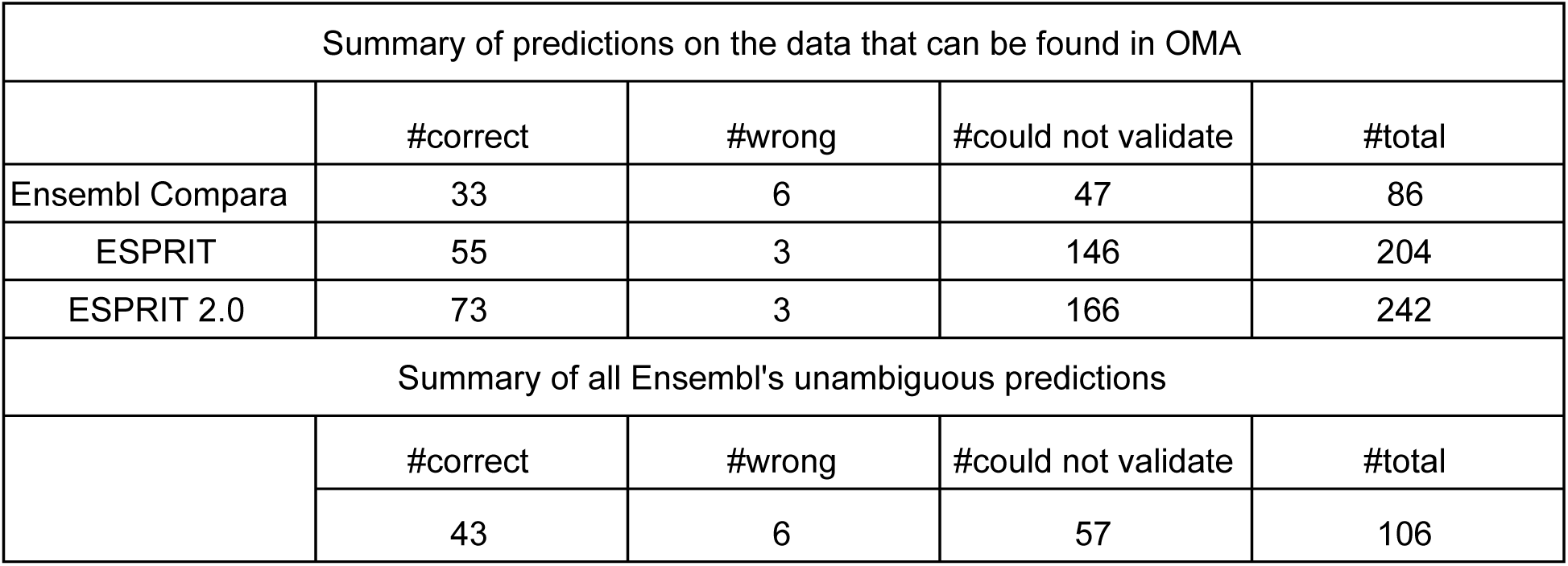
Comparison to Ensembl Compara and ESPRIT. Summary.

**Supplementary figure 4:**
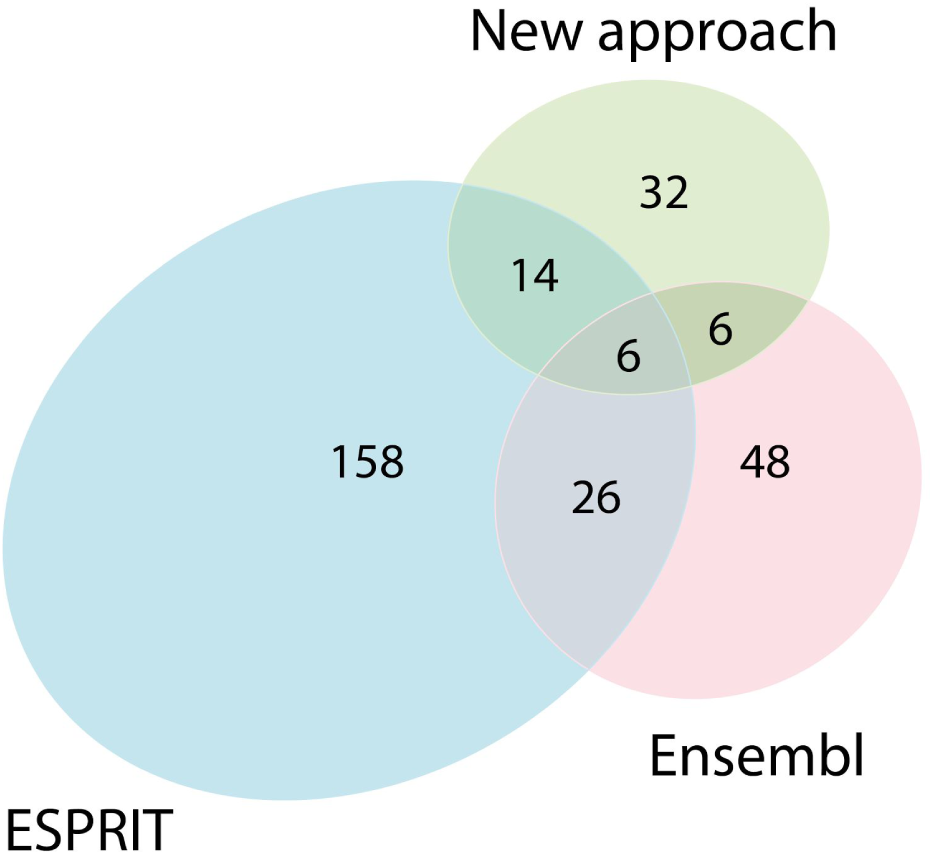
Comparison to Ensembl Compara and ESPRIT. The number of predictions inferred by each method on 3B survey sequence. The new approach combines Collapsing (threshold 0.95) and LRT (significance 0.01).

### Discussion

#### Mistakes in the likelihood ratio test

**Supplementary figure 5:**
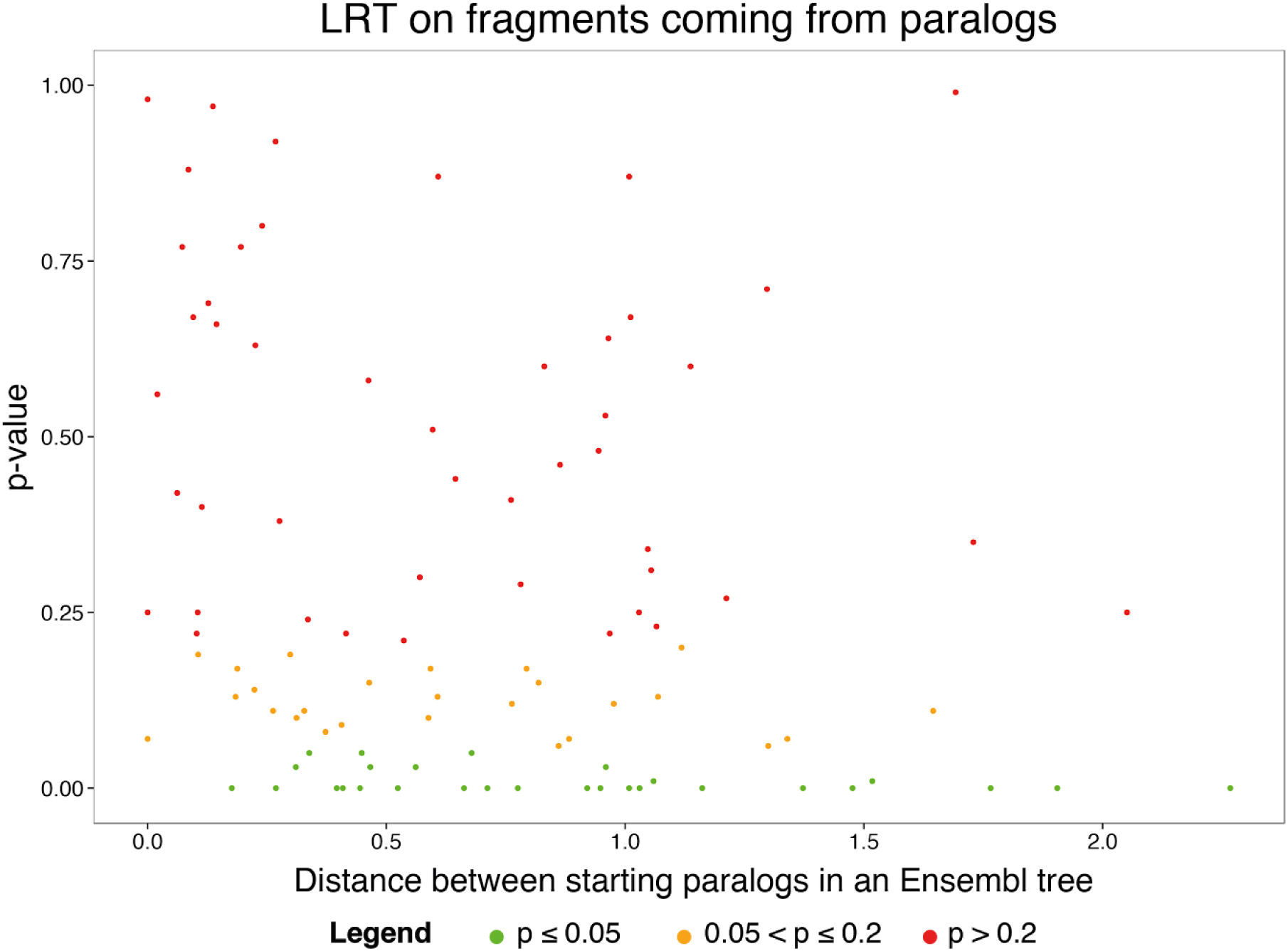
The relationship between paralog distance (expected number of changes per site) and p-value for the likelihood ratio test when applied to random fragments coming from the paralogs.

#### Fragments coming from regions that have evolved at different rates

Gene TRAES3BF091400260CFD_t1 was split at a random position. The p-value of the likelihood ratio test is probably low, i.e. in favour of the hypothesis that fragments come from paralogous sequences, due to dissimilar evolutionary rates. Collapsing approach correctly infers them as fragments of the same gene when used with any of the thresholds {0.65, 0.75, 0.85, 0.9, 0.95} (Suppl. table 4).

The gene family was obtained from Ensembl Plants, release 31, alignments performed by Mafft v7.164b (Suppl. File alignments.tar.gz) and trees built with FastTree v2.1.8 (Suppl. fig. 6).

**Supplementary table 4:** Information on a case where the likelihood ratio test distinguishes fragments coming from the same gene.

| Gene | P-value | #references in the MSA | Length of the MSA | Length (gene)<br>Length (fragment1)<br>Length (fragment2) | Results from collapsing approach |
| --- | --- | --- | --- | --- | --- |
| TRAES3BF091400260CFD_t1 | 0.02 | 3 | 355 | 244<br>92<br>152 | Split gene |
| Reference sequence |  |  | PAM distance (reference, fragment1) | PAM distance (reference, fragment2) |  |
| TRIUR3_21111-P1 |  |  | 43.06 | 734 |  |

**Supplementary figure 6:**
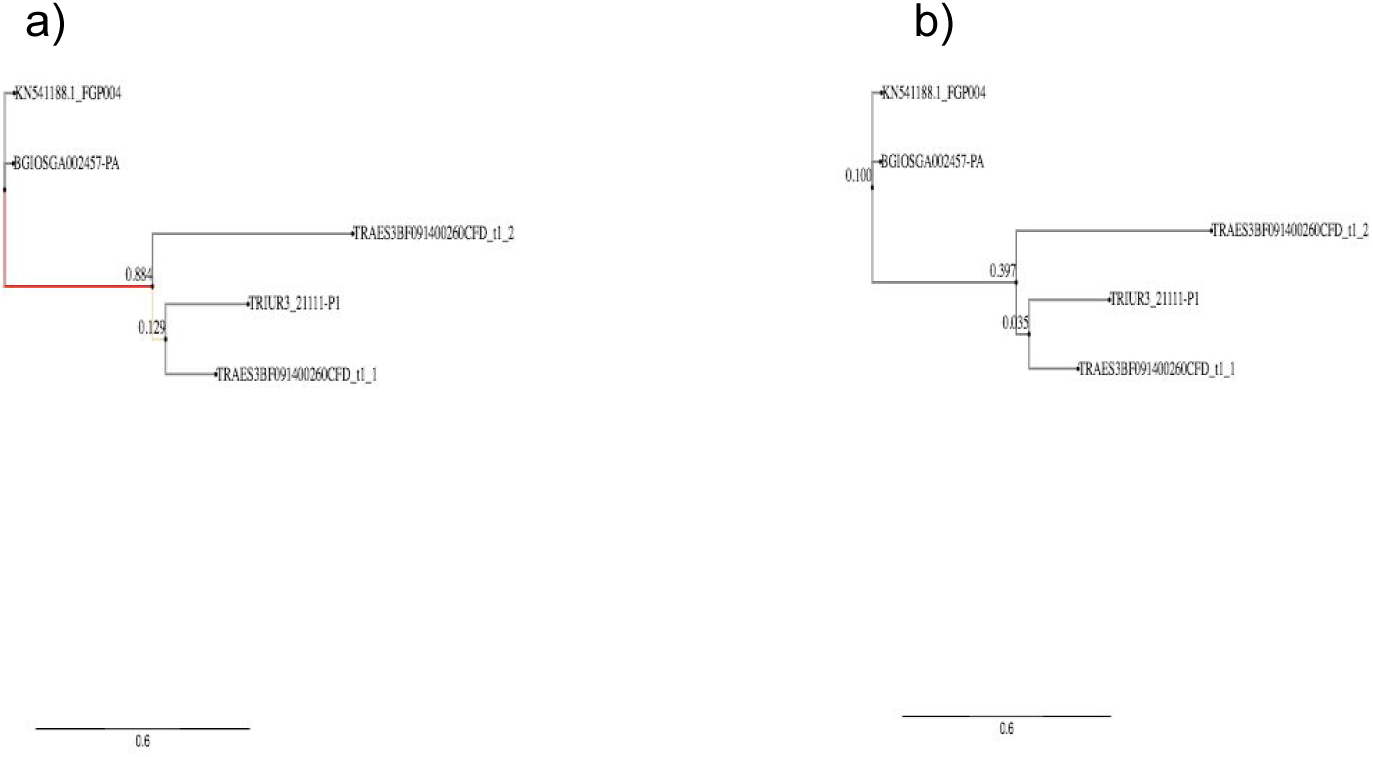
Gene tree containing fragments of the TRAES3BF091400260CFD_t1 gene. a) With branch bootstrap supports, b) With branch lengths.

#### Mistakes on distant sequences

To illustrate more cases where the likelihood ratio test makes incorrect inference, we also provide information on three cases (Suppl. table 5, alignments in Suppl. File esprit2_alignments.tar.gz). In all of them, we introduced fragmentation on distant paralogous sequences which should be sufficient to distinguish fragments as paralogous. Yet, based on the p-values and levels of significance of the test, we could not reject the hypothesis that fragments come from the same gene.

Given the size of the gene families, non-conserved long alignments and rather short candidate fragments, there is likely the lack of information contained in the alignments and fragments to make correct inference. However, there is enough information for the collapsing approach to infer that pairs in question are paralogous.

**Supplementary table 5:** Information on three cases where the likelihood ratio test fails to distinguish paralogous sequences.

| Gene1 | Gene2 | Distance_in_Ensembl_tree (gene1, gene2) | P-value | #genes in the MSA | Length of the MSA | Length (gene1)<br>Length (fragment 1) | Length (gene2)<br>Length (fragment2) | Results from collapsing approach |
| --- | --- | --- | --- | --- | --- | --- | --- | --- |
| TRAES3BF071900030CFD_t1 | TRAES3BF182800010CFD_t1 | 2.05 | 0.25 | 1040 | 11165 | 464<br>366 | 1160<br>83 | paralogs |
| TRAES3BF117700060CFD_t1 | TRAES3BF136700020CFD_t1 | 1.73 | 0.35 | 1293 | 16234 | 554<br>140 | 368<br>181 | paralogs |
| TRAES3BF075600050CFD_t1 | TRAES3BF171600010CFD_t1 | 1.69 | 0.99 | 1258 | 4924 | 243<br>81 | 207<br>109 | paralogs |

### Datasets

The datasets referenced in this document and the main body (simulated splits, comparison, predictions, case studies, etc.) can be downloaded from http://lab.dessimoz.org/17_esprit2.

## Footnotes

1 The OMA Browser release containing 3B survey sequence is older than the one containing 3B reference sequence. Hence, assemblies for some reference species differ between the releases as can be noticed from Suppl. table 1 and Suppl. table 2.

